# Enlighting the toxinological dark matter of spider venom enzymes

**DOI:** 10.1101/2024.02.27.582330

**Authors:** Josephine Dresler, Volker Herzig, Andreas Vilcinskas, Tim Lüddecke

## Abstract

Spiders produce highly adapted venoms featuring a complex mixture of biomolecules used mainly for hunting and defense. The most prominent components are peptidic neurotoxins, which have been the focus of research and drug development, whereas venom enzymes have been largely neglected. Nevertheless, investigation of venom enzymes not only reveals insights into their biological functions, but also provides templates for future industrial applications. Here we compared spider venom enzymes contained in the VenomZone database and in other publicly available proteo-transcriptomic datasets. We found extensive discrepancies between these sources, revealing a previously unrecognized abundance and diversity of venom enzymes. Furthermore, we assigned the reported enzymes to cellular processes and known venom functions, including toxicity, prey pre-digestion, venom preservation, venom component activation, and venom spreading factors. Our study reveals a gap between databases and publications in terms of enzyme coverage which impedes development of new applications based on the rich and diverse spectrum of enzymes contained in spider venom.

## 1. Introduction

Spiders (Araneae) are ancient arthropod assassins that inhabit virtually all terrestrial ecosystems [1]. During their > 300 million years of evolutionary history, they evolved into highly effective predators of insects and other arthropods [2]. Despite the wide range of ecosystems conquered, their principal body plan has remained largely unchanged and is shared by all extant species across the three spider infraorders (Mesothelae, Mygalomorphae and Araneomorphae) [3–5]. The astonishing success of spiders in terms of biodiversity, evolutionary age and ecological versatility, is to a large extent rooted in key adaptions and biomolecular innovations such as pheromone chemistry, silk production, and, in particular, venom [2,6–8].

Venoms are complex chemical cocktails that evolved convergently multiple times in the animal kingdom [9]. They are actively injected from one animal into another, where they disturb vital physiological processes and cause damage or even death [10]. The bioactive components of venoms are described as toxins and are mostly of proteinaceous nature [10]. The three major biological functions of venom are hunting, defense and intraspecific competition, but additional minor functions include chemical communication and reproduction among others [11].

Spiders are one of the oldest venomous lineages and also one of the most diverse, with ~51,000 described species [1]. This has led to the widespread recognition of spiders as the world’s most successful group of venomous animals, at least among terrestrial lineages [2]. The complexity of spider venom is also unmatched, featuring hundreds to thousands of different polypeptide components alone [12]. All spiders except the family Uloboridae are venomous, which means they offer an immense library of bioactive components. Based on extrapolations from the limited number of species studied thus far, it has been estimated that up to 10 million biomolecules can be isolated from all extant species [13].

Most known spider venom toxins are short disulfide-rich peptides with an inhibitor cysteine knot (ICK) motif [13]. These usually have molecular masses < 7 kDa and are potent modulators of a wide range of ion channels. Their biological function is primarily trophic, and they are used to mount neurochemical attacks on insect prey following injection via a cheliceral bite [2,14]. However, several of these ICK peptides also show promising activity against human ion channels involved in a wide range of neuropathologies. This bioactivity, paired with the extraordinary stability conferred by the ICK fold [15], means that spider venom peptides are potent drug leads for the treatment of neurological diseases, including epilepsy, chronic pain, and post-stroke brain damage [13,16–19]. Unsurprisingly, spider venom biodiscovery has focused on novel and potentially druggable ICKs and has typically involved bioactivity-guided screening, where crude venom is fractionated by chromatography and the fractions are analyzed by mass spectrometry to identify peptides with desired activities [20]. More recent venomics strategies combine proteomics and transcriptomics to shed light on the composition of venoms [21]. When combined with biotechnology, this approach provides access to all venom proteins and peptides in a given source [22]. A major insight from modern venomics is that spider venoms not only contain small ICK peptides but also an array of high-molecular-weight proteins, often with putative enzymatic activity [2,23]. Enzymes are complex proteins containing domains that catalyze chemical reactions. In general, the classification and nomenclature of enzymes depends on the catalyzed reaction and the substrates involved. The IUBMB enzyme classification system uses a four-component numerical system to classify enzymes based on Enzyme Commission (EC) numbers. Seven different enzyme classes are recognized: oxidoreductases (EC 1), transferases (EC 2), hydrolases (EC 3), lyases (EC 4), isomerases (EC 5), ligases (EC 6) and translocases (EC 7) [24]. In contrast to the frequent in-depth studies of neurotoxic peptides, the bioactivities of spider venom enzymes are almost entirely unknown. The only exceptions are sphingomyelinase D enzymes from sicariid spiders and a few neurotoxin-processing enzymes involved in toxin maturation [25–29]. However, in some araneomorph spiders, enzymes may be the major venom components, implying they serve important biological functions [30,31]. This growing body of evidence suggests that spider venoms contain a vast resource of undiscovered enzymes that can be thought of as toxinological dark matter. This is unfortunate because a better understanding of those molecules might provide further insights into the chemical and evolutionary ecology of spider venoms and could be employed as a valuable resource for translational research [32,33].

Here we set out to provide the first holistic, structural assessment of the enzymatic diversity within spider venoms. Specifically, we analyzed the content of a manually curated venom database and mined putative enzymes from modern venomics research. We linked all the enzymes identified in both approaches with taxonomic placements and potential functions. The wealth of data we analyzed shows that spider venoms are a rich source of enzymes and that the diversity of enzyme families rivals the number of known ICK families. Given the strong research bias towards ICK peptides and the recent advent of modern venomics, future studies should include the analysis of venom enzymes in more detail.

## 2. Material and Methods

### 2.1 Database search

To investigate the diversity of known spider venom enzymes in public databases, we selected the VenomZone database as our source [34]. VenomZone is a publicly available database that is manually curated and contains entries of venom components representing six different lineages (snakes, spiders, cnidaria, insects, scorpions and cone snails) that are pre-categorized into distinct toxin families. We screened all 1540 family-grouped entries for spiders and retrieved all proteins assigned to known enzyme families together with their chemical classifications as per EC numbering and the taxonomic information for each species. The entries including the original research paper in which they have been first described were collected in a master table, which is presented in supplementary table S1. In addition to that all species names have been adjusted to the current nomenclature as standing in the world spider catalog in February 2024 [1].

### 2.2 Literature search

To identify spider venom enzymes other than those contained in the VenomZone database, we performed a literature search using Pubmed and applied the following search terms: “spider”, “venom”, “venomics”, “transcriptomics”, “proteomics”. Given that transcriptomic data alone are prone to false-positive hits, we only considered works in which the transcriptomic hits had been verified using proteomics or other forms of mass spectrometry to ensure the highest possible stringency. The identified studies were analyzed for venom proteins assigned to known enzyme families, similar to the procedure used in the database search. All hits were collected in a master table (supplementary table S2) together with the taxonomic information, which has been adjusted to the current nomenclature as given in the world spider catalog [1].

### 2.3 Classification of the identified enzymes

To classify all identified enzymes, we assigned their EC numbers (supplementary table S1 and S2) using ENZYME [35], a nomenclature database that describes each characterized enzyme type associated with an EC number. It was not possible to track down specific EC numbers for all described enzymes, so we also assigned enzymes to classes and subclasses. We also identified one superfamily and a series of multidomain enzymes whose members belong to several enzyme classes and subclasses simultaneously. Finally, we included the cysteine-rich secretory proteins, antigen 5 and pathogenesis related protein 1 (CAP) superfamily and lectin glycoproteins because they sometimes exhibit enzymatic activity although they are generally not considered as enzymes *sensu strictu*.

## 3. Results

### 3.1 VenomZone

VenomZone is a manually curated database of venom components containing 49 protein families, 17 of which are enzymatic. We found that nine known enzyme families occur in spider venoms, representing two distinct enzyme classes (hydrolases and isomerases). We also considered the potentially enzymatic glycoproteins. The enzymes identified in each class and the number of identified proteins for each family are discussed below and are summarized in Table 1.

**Table 1:**
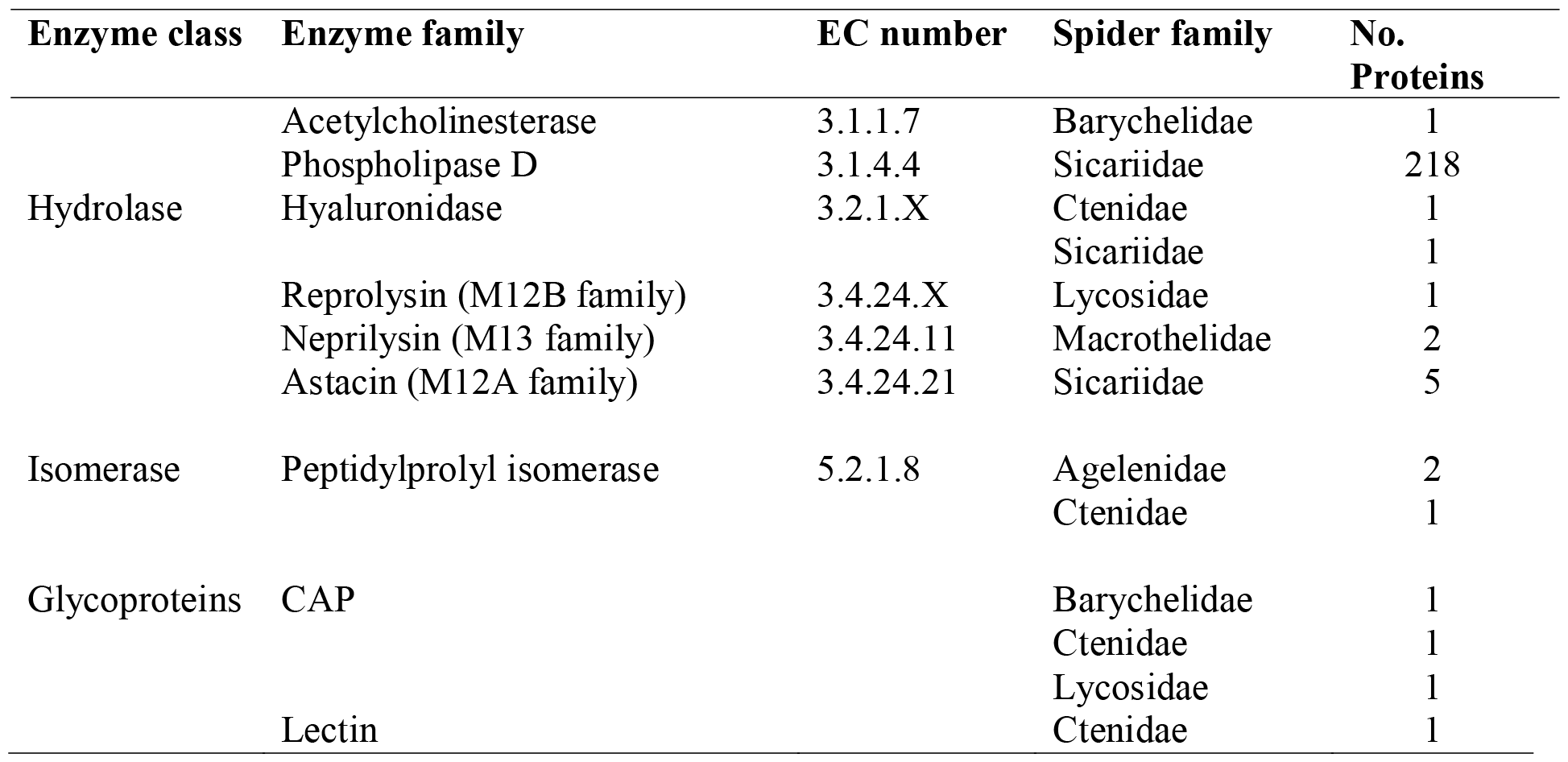
Diversity of spider venom enzymes present in the VenomZone database, showing assigned enzyme classes, families, EC numbers, taxonomic origins and number of proteins for each family.

#### 3.1.1 Hydrolases

Six distinct hydrolase families have been reported in spider venom. One acetylcholinesterase has been described in the brushed-foot trapdoor spider *Trittame loki* (family Barychelidae). The most diverse enzyme family is the arthropod dermonecrotic toxin (phospholipase D), also known as sphingomyelinase D. We found 218 representing the family Sicariidae, specifically 174 from the genus *Loxosceles*, 22 from the genus *Hexophthalma* and 22 from the genus *Sicarius*. Furthermore, two hyaluronidases have been reported, one each from *Phoneutria* (Ctenidae) and *Loxosceles* (Sicariidae).

All the remaining hydrolases were metalloproteases. Metalloproteases are a group of proteases that require an metal which is involved in their catalytic mechanism [36]. First, we manually classified two members of the neprilysin (M13 family) from the family Macrothelidae (genus *Macrothele*). These two members were not allocated to a specific protein family in VenomZone but we assigned them to the neprilysins based on sequence similarity. Second, we identified six members of the M12 family, comprising five astacins (M12A), all from the the genus *Loxosceles* in the family Sicariidae, and one partitagin (M12B, reprolysin) from the lycosid *Hippasa partita*.

#### 3.1.2 Isomerases

We identified three peptidylprolyl isomerases, two from Agelenidae (*Agelena*) and one from Ctenidae (*Phoneutria*).

#### 3.1.3 Glycoproteins

We identified four glycoproteins representing the cysteine-rich secretory proteins, antigen 5 and pathogenesis related protein 1 (CAP) family as well as one lectin. The CAP family is associated with diverse functions. CAPs isolated from snake venom are neurotoxic whereas those from hematophagous species are potentially hemotoxic [10], and little is known about spider venom CAPs. However, based on sequence similarity to *Conus* CAP proteins, they may represent a hitherto unrecognized lineage of proteases [30] and we therefore included them in our survey. One CAP has been described thus far from each of the families Barychelidae (*Trittame*), Lycosidae (*Lycosa*) and Ctenidae (*Phoneutria*). The single lectin we identified was presented in Ctenidae (*Phoneutria*). These carbohydrate-specific binding proteins that agglutinate cells or other materials are important for cell adhesion and defense [37,38].

### 3.2 Publications

To discover putative enzyme families that were previously overlooked, we collected all published spider venom proteo-transcriptomes from the last 19 years and analyzed them for the presence of enzymatic components. We identified 143 putative enzyme families that occur with varying numbers of identified protein for each enzyme family in spider venom (supplementary table S2). The enzymes represent seven distinct classes and are described in more detail below.

#### 3.2.1 Oxidoreductases

We identified 52 oxidoreductases belonging to 20 enzyme families, many from the spider family Theraphosidae. These include the synaptic vesicle membrane protein VAT-1 homolog-like, a thiol oxidase, two cysteine dioxygenases and two glucose-6-phosphate dehydrogenases in the genus *Acanthoscurria*, as well as 13 acyl-CoA oxidases and one uricase in *Chilobrachys guangxiensissis*. Furthermore, five phosphoglycerate dehydrogenases have been described, one from the Lamponidae (genus *Lampona*), one from the lycosid *Lycosa tarantula*, and the remaining three from the Theridiidae (genus *Latrodectus*). A gamma-interferon-inducible lysosomal thiol reductase has been described in the genera *Lampona* (Lamponidae), one in the *Acanthoscurria* and one in the Eresidae (*Stegodyphus*). A peroxidase has been described in the family Lamponidae (*Lampona*) and one in the family Eresidae (*Stegodyphus*) and dopamine β-hydroxylases have been identified in three spider families, one each in the Ctenidae (*Phoneutria*), the Lamponidae (*Lampona*) and the Theraphosidae (*Acanthoscurria*). One tyrosinase was identified in the theraphosid spider *Cyriopagopus schmidti*. Finally, a superoxide dismutase has been described twice in the family Ctenidae (genus *Phoneutria*), once in the family Philodromidae (genus *Tibellus*) and twice in the Theraphosidae (*Acanthoscurria*). Three retinal dehydrogenases have been reported in the Theraphosidae (*Acanthoscurria*) and one in the Eresidae (*Stegodyphus*). In the spider family Eresidae one glucose 1-dehydrogenase, one glutamate dehydrogenase and one glutathione peroxidase has been reported. A glyceraldehyde-3-phosphate dehydrogenase has been described in the Theraphosidae (*Acanthoscurria*) and in the Eresidae (*Stegodyphus*). The latter also featuring a 3-hydroxyisobutyrate dehydrogenase, a procollagen-lysine 5-dioxygenase and two thioredoxin-dependent peroxiredoxins.

#### 3.2.2 Transferases

We identified 49 transferases belonging to 27 enzyme families, spanning a range of diverse activities. Three methyltransferases have been described, one from the family Philodromidae (genus *Tibellus*) and two from the theridiid spider *Latrodectus tredecimguttatus*. The genus *Tibellus* also features one cullin-RING-type E3 NEDD8 transferase, one alanine transaminase, one aminoglycoside 2′′-phosphotransferase, one phosphoglycerate kinase, one trans-aconitate 2-methyltransferase and one type I protein arginine methyltransferase. Furthermore, two transketolase, one transaldolase, one arylamine *N*-acetyltransferase, one glycylpeptide *N*-tetradecanoyltransferase and one adenine phosphoribosyltransferase were identified in spiders from the family Theraphosidae. One polypeptide *N*-acetylgalactosaminyltransferase has been found in the genus *Lampona* (Lamponidae) along with a spermidine synthase. Three further spermidine synthases have been found in Theraphosidae (*Acanthoscurria*) along with three glutathione transferases, two from the genus *Chilobrachys* and one from the *Stegodyphus*. Additionally, one kynurenine-oxoglutarate transaminase, one protein kinase and one glycerate 3-kinase were reported in the theraphosid spider *Cyriopagopus schmidti*. Nine arginine kinases have been reported, two from the Linyphiidae (genus *Hylyphantes*), one from the Lycosidae (genus *Lycosa*), one from the Philodromidae (genus *Tibellus*) and five from the Theridiidae (genus *Latrodectus*). One nucleoside diphosphate kinase has been described in the genus *Lampona* and six non-specific serine/threonine protein kinases have been identified, four in the family Theraphosidae (two in the genus *Cyriopagopus*, one in *Pamphobeteus*, and one in *Acanthoscurria*) and two in the Theridiidae (genus *Latrodectus*). One galactosylceramide sulfotransferase and one alcohol sulfotransferase have been reported from the Eresidae (*Stegodyphus*) and one phosphatidylcholine-sterol O-acyltransferase from the Theraphosidae (*Acanthoscurria*). Finally, a myosin light chain kinase has been identified twice in the Theraphosidae (genus *Chilobrachys*) and a histidine kinase has been found in the genera *Tibellus* and *Chilobrachys*.

#### 3.2.3 Hydrolases

The most diverse enzyme class we identified in spider venom was the hydrolases, with 516 hydrolases belonging to 74 different enzyme families, comprising 21 acting on ester bonds, 12 glycosylases, 35 peptidases, five acting on carbon-nitrogen bonds and one acting on acid anhydrides, which are discussed in turn below.

The group acting on ester bonds includes three undetermined enzymes in the family Theridiidae, three undetermined nucleases, five undetermined phosphatases and one undetermined phospholipase. Others have been assigned more specifically. For example, two lipases have been reported in the Ctenidae (genus *Phoneutria*), one each in the Lamponidae (genus *Lampona*), the Theraphosidae (genus *Acanthoscurria)*, and the Theridiidae (genus *Parasteatoda*), and an unreported number of additional lipases has been found in the Theridiidae (genus *Steatoda*). The distribution of triacylglycerol lipases, phospholipases, acetylcholinesterases and other ester-specific hydrolases is summarized in supplementary table S3.

Among the glycosylases, we found amylases in the Tetragnathidae (*Tetragnatha*), the Theridiidae, the Trechaleidae (genus *Cupiennius*) and the Eresidae (*Stegodyphus*), as well as 25 hyaluronidases distributed among eight spider families. We also found one neutral alpha glucosidase (*Lampona*), one glycosyl hydrolase (*Cyriopagopus*) and 19 chitinases in seven spider families. Two lysozymes were identified, one each in the Araneidae (*Araneus*) and the Theridiidae (*Latrodectus*). We also found a α-galactosidase and two β-galactosidases in the Theraphosidae (*Acanthoscurria*) and three α-mannosidases, two in the Theridiidae (*Parasteatoda*) and one in the Eresidae (*Stegodyphus*). The distribution of these enzymes and other glycosylates is summarized in supplementary table S3.

The 35 peptidases we identified included cathepsins in five spider families and metalloproteases in the Ctenidae (*Phoneutria*) and the Theraphosidae (*Acanthoscurria*). Twenty-one undetermined peptidases were described in the family Theraphosidae (14 members, *Acanthoscurria*) and Theridiidae (seven members, *Latrodectus*). We identified ten angiotensin-converting enzymes in eight spider families. Carboxypeptidases were also widely represented, including two members of the carboxypeptidase C family in the Araneidae (*Argiope*) and Lamponidae (*Lampona*), one carboxypeptidase D each in the Theridiidae (*Parasteatoda)*, the Eresidae (*Stegodyphus*) and the Theraphosidae (*Acanthoscurria*), three members of the carboxypeptidase A family two in the Theraphosidae (*Acanthoscurria*) and one in the Trechaleidae (*Cupiennius*), two carboxypeptidases B from Theridiidae (*Parasteatoda*) and Eresidae (*Stegodyphus*), one carboxypeptidase E from Theraphosidae (*Acanthoscurria*) and two carboxypeptidases M from Tetragnathidae (*Tetragnatha*). We found 27 S1 proteases in ten spider families, and eight proprotein convertases in six spider families. Neprilysin was represented 198 times in eight spider families, 154 from the Atractidae (*Hadronyche*). We found 27 astacins in five families and an undefined number of further examples in the Theridiidae (*Steatoda*). The distribution of these enzymes and other glycosylates is summarized in supplementary table S3.

The five enzyme families specific for carbon-nitrogen bonds comprised an arylformamidase identified twice in the Philodromidae (*Tibellus*) and once in the Theraphosidae (*Cyriopagopus*), a ceramidase found once each in the Lamponidae (*Lampona*) and the Theraphosidae (*Acanthoscurria*), a pantetheine hydrolase in the Theraphosidae (*Acanthoscurria*) and the Eresidae (*Stegodyphus*), two dihydropyrimidinases from the Theraphosidae (*Chilobrachys*) and one 2-iminobutanoate/2-iminopropanoate deaminase from Philodromidae (*Tibellus*). The last of the hydrolases is the helicase which acts on acid anhydrides and has been detected in the Lamponidae (*Lampona*).

#### 3.2.3 Lyases

We identified 12 lyases belonging to five enzyme families. These comprised one aromatic L-amino-acid decarboxylase in the Araneidae (*Araneus*) and eight carbonic anhydrases, two in the Lamponidae (*Lampona*) and six in the Theraphosidae (*Acanthoscurria*), as well as one enolase in the Lamponidae (*Lampona*), one aconitate hydrolase in the Philodromidae (*Tibellus*) and one dermonecrotic protein 1 LiDl in the Sicariidae (*Loxosceles*).

#### 3.2.4 Isomerases

We identified 15 isomerases to four enzyme families. Six peptidyl-prolyl cis-trans isomerases have been reported, two each in the spider families Lamponidae (genus *Lampona*), Philodromidae (genus *Tibellus*) and Ctenidae (genus *Phoneutria*). Two triose-phosphate isomerases have been described from the Theridiidae (genus *Latrodectus*) as well as six protein disulfide isomerases, one from the Trechaleidae (genus *Cupiennius*), three from the Philodromidae (genus *Tibellus*), one from the Eresidae (*Stegodyphus*) and one from the Theraphosidae (*Acanthoscurria*). Finally, a single phosphomannomutase has been identified in the theraphosid spider *Cyriopagopus schmidti*.

#### 3.2.5 Ligases

We identified three ligases belonging to three enzyme families, only one of which could not be assigned a specific EC number. One ligase has been reported in the Theraphosidae (genus *Cyriopagopus*), one acyl-CoA synthetase from the araneid spider *Araneus ventricosus*, and one carbamoyl-phosphate synthase from the Theraphosidae (genus *Cyriopagopus*).

#### 3.2.6 Translocases

We identified 40 translocases belonging to two enzyme families. This includes 30 ATP synthases that have been reported in five spider families. One is from the family Lamponidae (genus *Lampona*), six from the Philodromidae (genus *Tibellus*), two from the Theraphosidae (one from genus *Chilobrachys* and one from *Cyriopagopus*), 19 from the Theridiidae (genus *Latrodectus*) and one from the Eresidae (*Stegodyphus*). Furthermore 10 Ca^2+^-transporting ATPases have been identified in the theridiid spider *Latrodectus tredecimguttatus*.

#### 3.2.7 Multidomain enzymes, glycoproteins and others

We identified 13 multidomain enzymes belonging to 5 enzyme families. For example, peptidylglycine monooxygenases contain two separate catalytic domains, but the enzyme is assigned only one EC number [39]. Eight have been described in four spider families, four in the Theraphosidae (genus *Acanthoscurria*), one in the Theridiidae (one genus *Parasteatoda*), one in the Trechaleidae (genus *Cupiennius*), two in the Eresidae (*Stegadyphus*) and more in the theridiid genus *Steadoa* but the precise number has not been reported. One α-aminoadipic semialdehyde synthase has been detected in the philodromid spider *Tibellus oblongus* and one histone-lysine *N*-methyltransferase ASH1L from the Theraphosidae (*Acanthoscurria*). Two cyclin-dependent kinases have been found in the Philodromidae (genus *Tibellus*) and one dual-specificity phosphatase 23 in the Theraphosidae (genus *Cyriopagopus*). In contrast to the peptidylglycine monooxygenase, these enzymes have more than one EC number, each representing a different enzymatically active domain.

We identified 78 glycoproteins belonging to 2 enzyme families, i.e.73 CAPs and 5 lectins distributed throughout the spider phylogenetic tree. Three CAPs were reported in the Atracidae (*Hadronyche*), 26 in the Araneidae (twelve genus *Argiope* and 14 genus *Araneus*), one in the Barychelidae (*Trittame*), five in the Ctenidae (genus *Phoneutria*), three in the Linyphiidae (genus *Hylyphantes*), four in the Lycosidae (*Lycosa*), two in the Philodromidae (*Tibellus*), two in the Pholcidae (genus *Physocyclus*), two in the Tetragnathidae (genus *Tetragnatha*), 14 in the Theraphosidae (nine genus *Acanthoscurria*, five genus *Pamphobeteus*), nine in the Theridiidae (one genus *Parasteatoda*, eight genus *Steatoda*) and two in the Trechaleidae (*Cupiennius*). Lectin has been found once in the families Ctenidae (*Phoneutria*) and Lamponidae (*Lampona*) and twice in the Theraphosidae (*Acanthoscurria*).

One superfamily containing α/β-hydrolases has been detected in spider venom. One member of this superfamily has been identified in the spider *Cyriopagopus schmidti* (Theraphosidae). Given that the superfamily contains enzymes representing enzyme classes EC 1–6, no further classification was possible.

### 3.3 Discrepancies between database entries and venomics studies

A comparison of enzymes deposited in databases and those mined from venomic data revealed extensive discrepancies. Only nine enzyme families are described in VenomZone, whereas we identified 143 enzyme families by screening published venomic data. Eight of these enzyme families were found in both sources, revealing that 135 enzyme families identified in the literature are not found in databases. Only the partitagins (M12B family) are exclusively found in VenomZone, and even this family may be included among the many undetermined metalloproteases described in the literature that are not specifically identified in the proteo-transcriptomics data.

### 3.4 Biological function of the enzymes

Spiders seem to utilise all distinct enzyme classes with the majority being hydolases, transferases and oxidoreductases (Figure 1). To develop a deeper understanding of the identified enzymes we sought to assign putative venom functions using the functional classifications previously established for arachnid venom enzymes, principally the classification of enzymes detected in scorpion venom [40] and spider venom [41].

**Figure 1:**
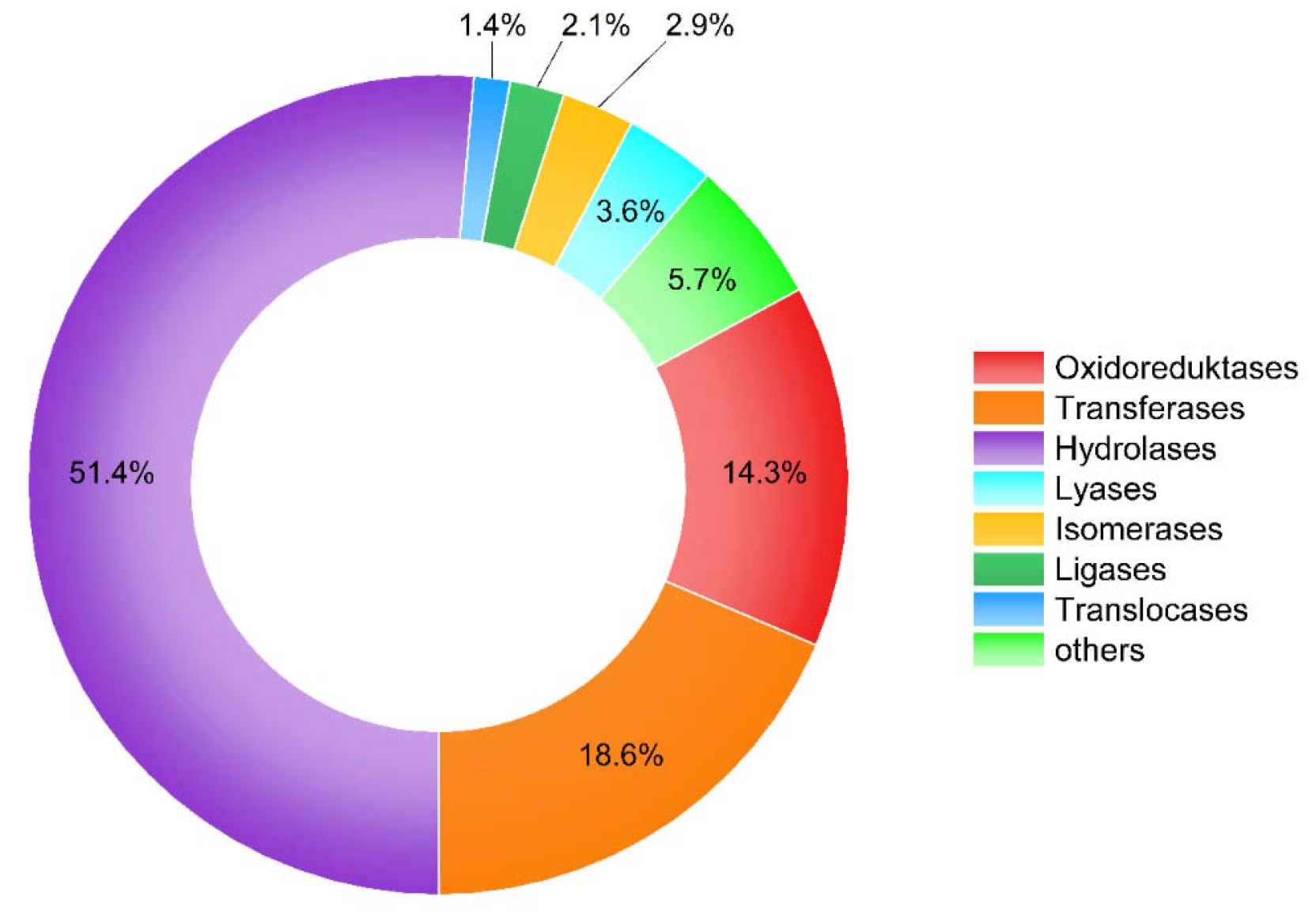
The diversity of spider venom enzymes. Relative distribution of spider venom enzyme families from VenomZone entries and proteo-transcriptomic studies per enzyme class. The category “others” includes glycoproteins, multidomain enzymes and the superfamily.

According to the Delgado-Prudencio classification [40], we were able to assign 25 of the 144 enzymes to physiological functions. Members of the phospholipase D and acetylcholinesterase families were assigned as toxic enzymes, whereas triacylglycerol lipases, chitinases, amylases, alpha-galactosidases and ceramidases are thought to be involved in the pre-digestion of prey. Furthermore, 5′ nucleosidases, hyaluronidases, angiotensin converting enzymes and coagulation factor Xa may act as spreading factors, whereas dopamine β-monooxygenase, lysozymes, carboxypeptidases E and peptidylglycine monooxygenases are thought to activate venom components. Peroxidases, superoxide dismutases and carbonic anhydrases putatively have preservative functions whereas thioredoxin-dependent peroxiredoxins, phospholipases A2, lipases, carboxypeptidases B, trypsin, neprilysin and astacin have multiple enzymatic functions in venom (supplementary table S4). Figure 2 shows enzyme classes linked to physiological functions. Despite the remarkable abundance of hydrolases, only some oxidoreductases and one lyase seem to have known associated venom function. Hydrolases play a role in every function of venom except preservation. Interestingly, the functions assigned to oxidoreductases and the one lyase include preservation, with oxidoreductases also activating venom components.

**Figure 2:**
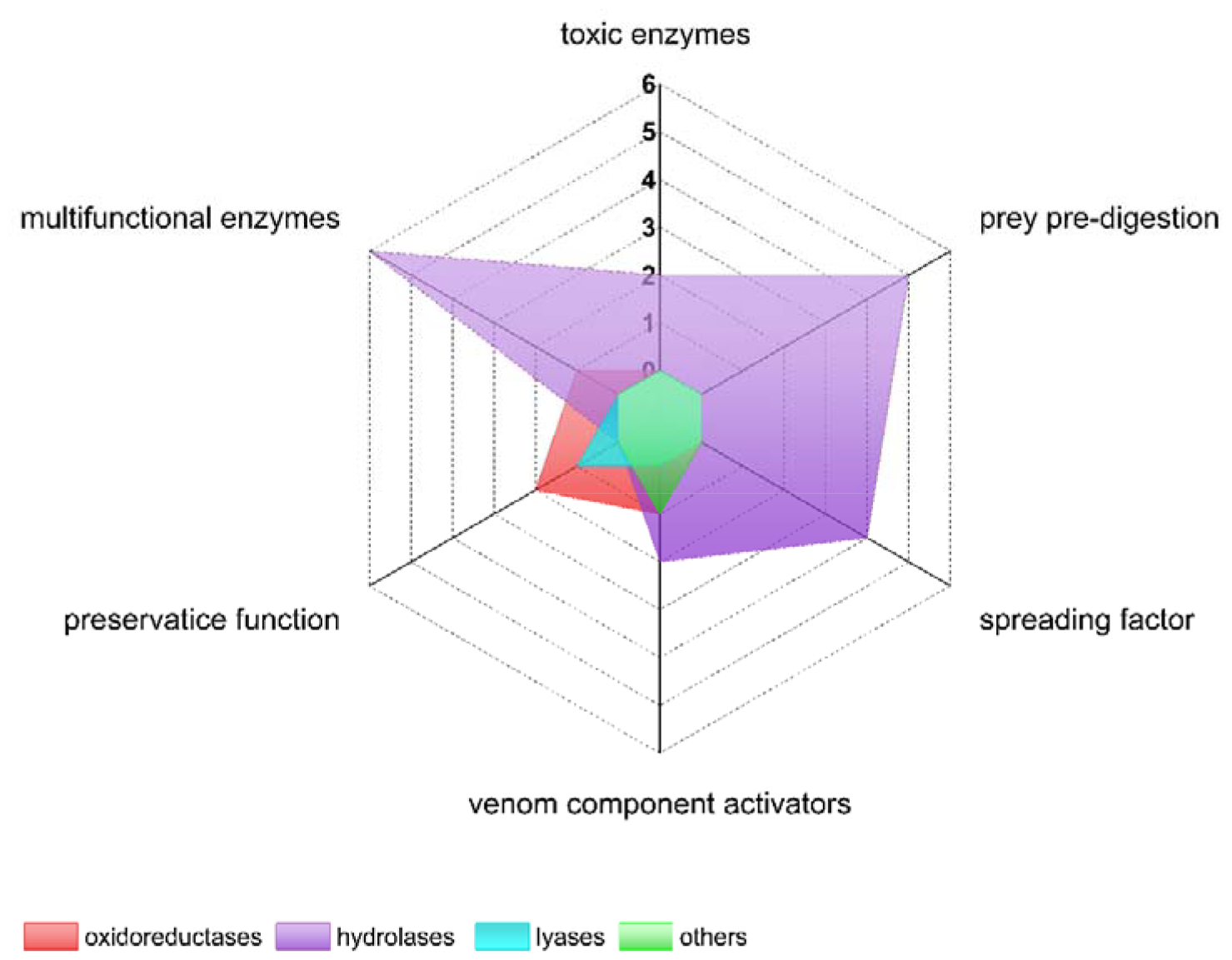
The spectrum of biological functions of spider venom enzymes. The radar plot shows the number of herein identified spider venom enzymes classes for each of the previously established functional classes of arachnid venom enzymes. The category “others” includes glycoproteins, multidomain enzymes and the superfamily. Counts show the number of enzymes per category.

We also assigned 13 of the enzymes to cellular functions such as metabolism or a contribution to cellular components [41]. Five of them are oxidoreductases (glucose-6-phosphate dehydrogenase, glucose 1-dehydrogenase, γ-interferon-inducible lysosomal thiol reductase, and dopamine β-monooxygenase, glutathione peroxidase), two are transferases (arginine kinase and nucleoside diphosphate kinase), five are hydrolases (α-mannosidase, hexosaminidase, proprotein convertase 1, S1 proteases and metalloendopeptidases), one is a lyase (carbonic anhydrase), two are isomerases (peptidylprolyl isomerase and protein disulfide isomerase), one is the multidomain enzyme peptidylglycine monooxygenase and lectins (supplementary table S4).

## 4. Discussion

### 4.1 The unrecognized diversity of spider venom enzymes

In this study, we have provided the first systematic analysis of the distribution and abundance of spider venom enzymes. Our results show many different enzymes spread across the spider tree of life. Overall, 144 enzyme families have been described from 18 spider families, 9 in the VenomZone database whereas 135 are exclusively found in proteo-transcriptome data. These are distributed in all enzyme classes, which highlights the chemical diversity of spider venoms (Figure 1). Neurotoxins are well-known as the major component of spider venom whereas enzymes are considered to be minor components [25–29]. Because of this, and given their potential as drug leads, neurotoxins have been the focus of many studies, whereas enzymes have been mostly overlooked. We found that venom enzymes show remarkable diversity, comparable to that of the neurotoxins. In some cases, the number of sequences identified in the proteome data was also comparably high. For example, *Phoneutria nigriventer* venom proteome consists of ~42% neurotoxins and ~43% enzymes [41] whereas the venom of *Steatoda nobilis* only features ~15% enzymes compared to ~49% toxins [42]. The abundance of the identified spider venom enzyme sequences therefore seems to vary depending on the species and further adds to the overall diversity of spider venom composition.

Spiders are mainly investigated if they are large, like many of the mygalomorphs [19,20], or if they are medically relevant in humans, such species in the genera *Loxosceles* or *Latrodectus* [43]. Considering the number of enzymes described throughout the phylogenetic tree, we see the greatest abundance in modern spiders from the infraorder Araneomorphae (Araneoidea and RTA clade) and in the more primitive family Theraphosidae which is part of the Mygalomorphae (Figure 3). This is supported by the number proteo-transcriptomic studies investigating theraphosid (8), theridiid (5), araneid (3) and ctenid (3) venom. Theraphosid spiders are among the largest spiders, explaining the increased attention they receive from the scientific community. A large proportion of reported toxins are derived from mygalomorph species [44], while other spider families have been largely overlooked in venom research [43]. However, we found that the situation is currently undergoing a change with recent publications focusing on four previously neglected spider families [45].

**Figure 3:**
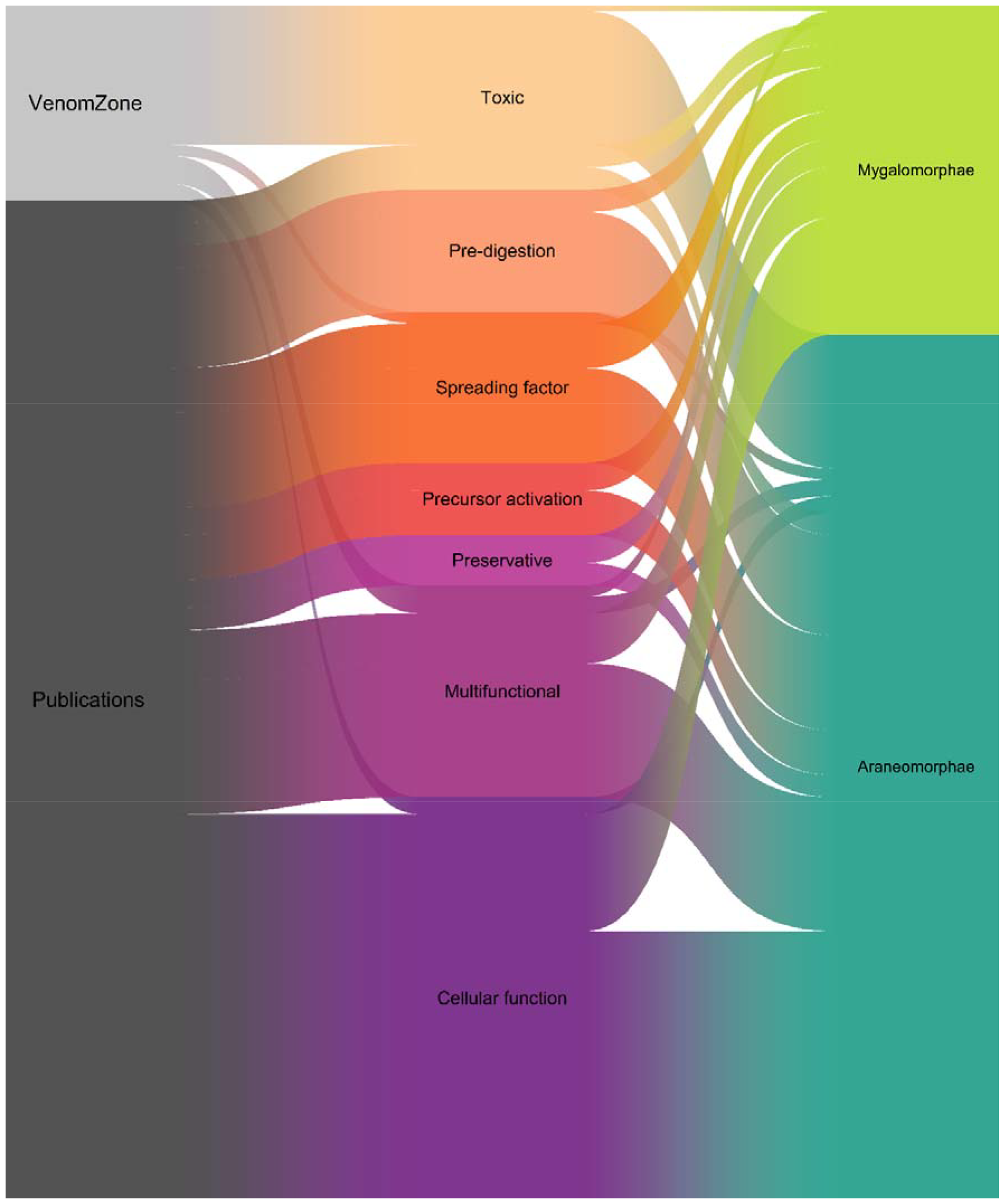
Identified spider venom enzymes in light of biological function and taxonomic origin. Sankey plot showing the relationship between venom enzymes identified from the VenomZone database and publications (left side), their associated functions (middle part) and presence in spider infraorders (right side).

### 4.2 Consequences of the absence of spider venom enzymes from databases

Our study revealed a large difference in the distribution of spider venom enzymes in databases and publicly available data. We found that the number of identified spider venom enzymes increased in the recent past, including many small araneomorph species, but these data have apparently not yet found their way into the respective databases. The traditional venom characterization pipeline involves bioactivity screening of crude venoms, which is dependent on the often limited venom yield of the chosen source species [44]. More recently, the characterization of venom has been facilitated by omics technologies particularly transcriptomics and proteomics. These combine bioinformatics tools with existing functional annotations of known toxins from databases to identify proteinaceous components and predict their function from minuscule starting material [44,46–48]. This enables the examination of venoms that were traditionally not accessible [44]. The apparent absence of most spider venom enzymes from public databases may therefore reflect the dominance of components identified using the less sensitive methodologies, the restricted range of target species previously examined, and the persistent focus on small neurotoxic peptides from spider venoms. Another explanation could be that public databases have not yet caught up with more recent discoveries in spider venom enzymes.

On another note, the widespread neglect of enzymatic components in venom databases has potential effects for future biodiscovery programs. In a venomics experiment, lineage specific databases are the principal means to functionally annotate amino acid sequences encoding for venom components. If suitable database entries for a given protein class are not present in the chosen database, then functional annotations will not be feasible. Our analysis shows that a vast majority of enzyme families that have been identified from spider venoms are absent from the VenomZone database. Thus, if only VenomZone is used for venomics experimenta, the conducted venom profiling will be blind towards the missing components. Therefore, the absence of spider venom enzymes from databases bears the risk of resulting in incomplete venom profiling that lacks important biological and translational details. Our findings underpin the importance of well curated taxon-specific databases for toxinological research and suggest that more concerted efforts are required to include hitherto overlooked venom components. This might not only lead to important discoveries revolving around spider venom biology, but also yield a plethora of useful enzymes for biotechnological applications.

### 4.3 Major patterns among spider venom enzymes and their biological functions

Spiders use their venom mainly for hunting and/or defense, and thus contain a complex mixture of different components to support these efforts [2]. In order to gather a better understanding of the functional role of enzymes across spider venoms, we classified all identified enzymes towards their putative function according to previously established categories. Considering the classifications proposed by Delgado-Prudencio et al. [40] and Diniz et al [41] we were able to assign 25 of 144 enzyme families to different venom functions and 17 enzyme families to cellular functions. In addition to this classification, we further classified some enzyme families based on their catalyzed reaction or given their previously determined function (supplementary table S3) [10,34,49,50]. We were able to assign eight additional enzyme families to putative venom functions, seven potential venom component activators, two potential prey pre-digestion enzyme and four potential cellular function enzymes.

Cellular functions relate to the production, maturation and secretion of complex toxin components include transcription, protein folding, transport and post-translational modifications, as well as ubiquitination and proteolysis [50]. The Delgado-Prudencio classification describes six global venom functions based on physiological effects in relation to the enzymatic activities and their possible role in envenomation: toxic enzymes, prey pre-digestion enzymes, spreading factors, venom component activators, preservatives, and multifunctional enzymes [40]. Toxic enzymes are one of the most relevant enzymatic venom compounds because they are directly related to the toxicity and physiological effects triggered by envenomation [40]. Enzymes with digestive functions are important for prey pre-digestion, whereas enzymes that compromise tissue integrity help to promote the distribution of other venom components [40]. Enzymes are also required to activate venom components by proteolysis or other forms of modification [40]. Enzymes with preservative functions help to eliminate reactive oxygen species (ROS), which are the intermediates or end products of cellular metabolism [51]. Venom components are susceptible to ROS, which results in a shorter half-life [52]. Many venom enzymes have multiple functions, and these have been assigned to a “multifunctional enzyme” category rather than specifying the individual functions [40].

Oxidoreductases are required for toxin folding. Their diversity in spider venom indicates that the venom gland is highly active, producing many venom peptides/proteins that require assistance to adopt their correct tertiary structures [41]. This enzyme class is the third most common in our study. Some have been assigned a preservative function, while others are required to activate venom components. Two examples for the protection against oxidative stress are superoxide dismutases and peroxidases. Whereas superoxide dismutases are associated with protection against oxidative stress in insects [53], peroxidases are also presumed to contribute to this function [40].

Transferases modify other proteins by transferring functional groups. For example, kinases in endoparasite venom chemically modify proteins by adding phosphate groups that help to inhibit the host immune system [54]. We found many transferases, mostly kinases, but we were unable to assign specific venom functions. Because they contribute to the inhibition of the host immune system in endoparasites, they may act as defense toxins in spider venom by suppressing the prey’s immune system during envenomation [55]. Additionally, the Diniz classification assigned some kinases to cellular functions [41], nucleoside diphosphate kinases for example primary participate in metabolic processes [50]. The injection of *Loxosceles intermedia* crude venom into rats increased the plasma levels of several transferases, and induced histopathological changes that indicated hepatic lesions [56]. Transferases may therefore play a role in the inhibition of the prey’s immune system, or may fulfil cellular functions beyond the function of venom, or may have a direct physiological effect in the prey. Hydrolases break down larger biomolecules into units that can be assimilated by the digestive system [57] and facilitate the metabolism of extracellular matrix components [58– 60]. The degradation of biomolecular components not only helps with pre-digestion but also facilitates the diffusion/spreading of other venom components, which explains why many of these enzymes have been assigned as pre-digestive or spreading factors. Many hydrolases in animal venoms are metalloproteases or serine proteases, which participate in the harmful effects of snake (Elapidae, Viperidae) and spider (Sicariidae) venoms [61,62]. Phospholipase A2 and acetylcholinesterase are hydrolases that have been assigned as toxic enzymes. Acetylcholinesterases cause tissue lesions or neurological alterations, and are responsible for the flaccid paralysis triggered by snake venom [63]. Protein disulfide isomerases, protein tyrosine phosphatases and trypsin have been associated to cellular functions by supporting the protein folding or post-translation modifying venom components [50]. Hydrolases may therefore be involved in the pre-digestion, as spreading factors as well as being part of cellular functions.

## 5. Conclusion and future perspectives

We present the first systematic and comprehensive analysis of enzyme diversity in spider venoms based on database searches and manually screening previously published proteo-transcriptomic datasets. While enzymes were historically considered insignificant components of spider venoms, we demonstrated that this is misleading because it masks the fact that enzyme diversity is comparable to that of neurotoxins. We assembled 144 enzyme families and proteins with putative enzymatic activities representing 18 spider families. Most of the enzymes were hydrolases, oxidoreductases or translocases. We were able to assign 43 enzyme families to different venom functions or associated cellular functions. The venom functions included venom toxicity, spreading factors, venom activation and preservation, or prey pre-digestion, whereas the cellular functions included metabolism and the formation of cellular components. Most of the described enzymes were not previously associated with specific catalytic activities or venom functions. While theoretically annotating enzymes to specific functions is an important first step, extensive future experiments are required to experimentally validate these functions, which then helps in further improving theoretical annotations.

## 6. Acknowledgements

We would like to thank the authors of “The enzymatic core of scorpion venoms” for being an inspiration, as well as Richard M. Twyman for commenting and editing this manuscript.

